# Competition, Mutualism, and Host Immune Control in a Cancer Microbiome

**DOI:** 10.1101/2025.05.06.652496

**Authors:** Eeman Abbasi, Ammal Abbasi, Erol Akçay

## Abstract

The microbiome functions as an ecological community, where diverse microbes engage in metabolically mediated interactions such as mutualism and competition. Host immune response can regulate microbial community richness and abundance, which in turn can shape the prevalence of different ecological interactions within the microbiome. Theory predicts that host immune states shift dominant interaction modes among microbes: inflammation favors competition, while immunosuppression favors mutualism. We test these theoretical predictions using the stomach cancer microbiome data through integrated genomic and metabolic analyses. We observe that tumors with high-richness and high-abundance microbiomes were associated with increased mutualistic interactions, whereas tumors with low-richness and low-abundance microbiomes had fewer mutualistic interactions. Host immune gene expression in the high-richness and abundance group was suggestive of a dysregulated or immunosuppressed tumor microenvironment, whereas in the low-richness and abundance group immune signatures were indicative of intact immune function. Notably, competitive interactions remained relatively consistent between groups, whereas mutualism varied markedly, highlighting its sensitivity to shifts in immune state. Finally, the microbiome and host immune states were linked to patient clinical outcomes, with high-richness and abundance microbiomes associated with poorer survival and elevated expression of immune markers linked to adverse prognosis. These results reveal how host immune control can covary with ecological interactions within the microbiome, and the potential consequences of these interactions for host health.

## 1 Introduction

The microbiome encompasses a diverse array of microorganisms, including bacteria, fungi, viruses, and archaea. Their significance in host health and potential as therapeutic targets is increasingly recognized. While the microbiome can promote host well-being, it can also act as a catalyst or indicator of disease onset, posing a challenge in determining causality (Hou et al., 2022). Therefore, understanding how the mechanisms that shape microbial communities and how they interact with hosts is crucial.

Recently, the role of ecological interactions in shaping microbial community dynamics and their implications for hosts has garnered significant attention (Coyte & Rakoff-Nahoum, 2019; Coyte et al., 2015; Goyal et al., 2021; Levy & Borenstein, 2013). The microbiome operates much like an ecological community, where various microbes interact within the ever-changing host environment (Gilbert & Lynch, 2019). Many of these interactions are metabolic, based on breaking down and utilizing different resources in the host environment (Zaneveld et al., 2017). Each microbe possesses a collection of genes encoding enzymes that break down large, complex macro-molecules (Martens et al., 2011). Because no single microbe can fully metabolize complex molecules, metabolic contribution from multiple microbes is required for complete digestion (Zengler & Zaramela, 2018). Moreover, microbial metabolism is often characterized by the leakage of metabolic intermediaries and by-products into the extracellular environment. The leakage of metabolites into the extracellular environment and the successive digestion of compounds by multiple microbes results in metabolic interdependencies fostering cross-feeding interactions. These interactions can become mutualistic if the species involved receive a two-way benefit (Hesse & O’brien, 2024). However, there are instances where microbes overlap in their need for the same metabolite for growth, leading to competitive interactions. Previous studies suggest that competition is the dominant interaction among culturable bacteria in laboratory settings (Foster & Bell, 2012). In contrast, positive interactions are more prevalent in host-associated living systems compared to free-living ones (Machado et al., 2021). These differences in the prevalence of a specific interaction observed between labgrown and host-associated microbial communities highlight the importance of the host environment. Host control of oxygen and nutrient gradients, diverse spatial topography and the immune cell populations can collectively affect the structure of the microbial community (Gralka et al., 2020). Hence the landscape of the ecological interactions observed for a microbial community is an emergent property that relies not just on the microbes present but also the surrounding environment.

In addition to the ecological interactions within it, it is well established that microbiomes are fundamentally structured by mechanisms of host control, such as nutrient provision (Mansour et al., 2021) or immune responses (Belkaid & Hand, 2014). These host control mechanisms can also shape and interact with the community level ecological interactions that we observe within the microbiome. However, there is very little known about this process.

Microbiomes and host immune responses to them are a classic example of coevolution. Both the innate and the adaptive immune response in vertebrates interact in tandem to maintain the commensal relationship with the resident microbiota while eliminating any pathogens (Thaiss et al., 2016). The innate immune response detects any foreign molecular patterns through pathogen recognition receptors and concurrently coordinates with the adaptive immune response to generate and identify antigens to combat foreign invaders through antibody-mediated mechanisms (Belkaid & Hand, 2014; Thaiss et al., 2016). In a state of host homeostasis, the host maintains a diverse microbiome (Zaneveld et al., 2017), accompanied by the balanced immune population exhibiting both inflammatory and regulatory immune markers (Belkaid & Hand, 2014). Disruptions to this homeostatic state can have consequences for the microbial community, leading to differential microbial community structure observed during the state of inflammation or immunosuppression.

Previous theoretical work (Abbasi & Akçay, 2022) suggests that when the host’s innate immune system exerts tighter control over the microbiome (such as in an inflammation state), this suppresses community abundance and renders the community more susceptible to external invasions indicating that a tighter leash suppresses mutualistic microbial communities. This is because mutualistic communities reach high abundances by relying on other species for resources and thereby resist invasion. However, competitive communities appear to be less affected by the inflammatory state of the host as they rely on the external environment and not on other species for resources to support growth and can reduce the population sizes of other species through competitive exclusion (see Figure 1). Conversely, in an immuno-suppressive state, mutualistic communities flourish, fostering greater species diversity and abundance (see Figure 1). Building on these theoretical insights, we propose that communities exhibiting high richness and abundance would demonstrate elevated levels of mutualistic interactions, in comparison to communities with lower richness and abundance and would have less stringent host immune regulation.

**Figure 1:**
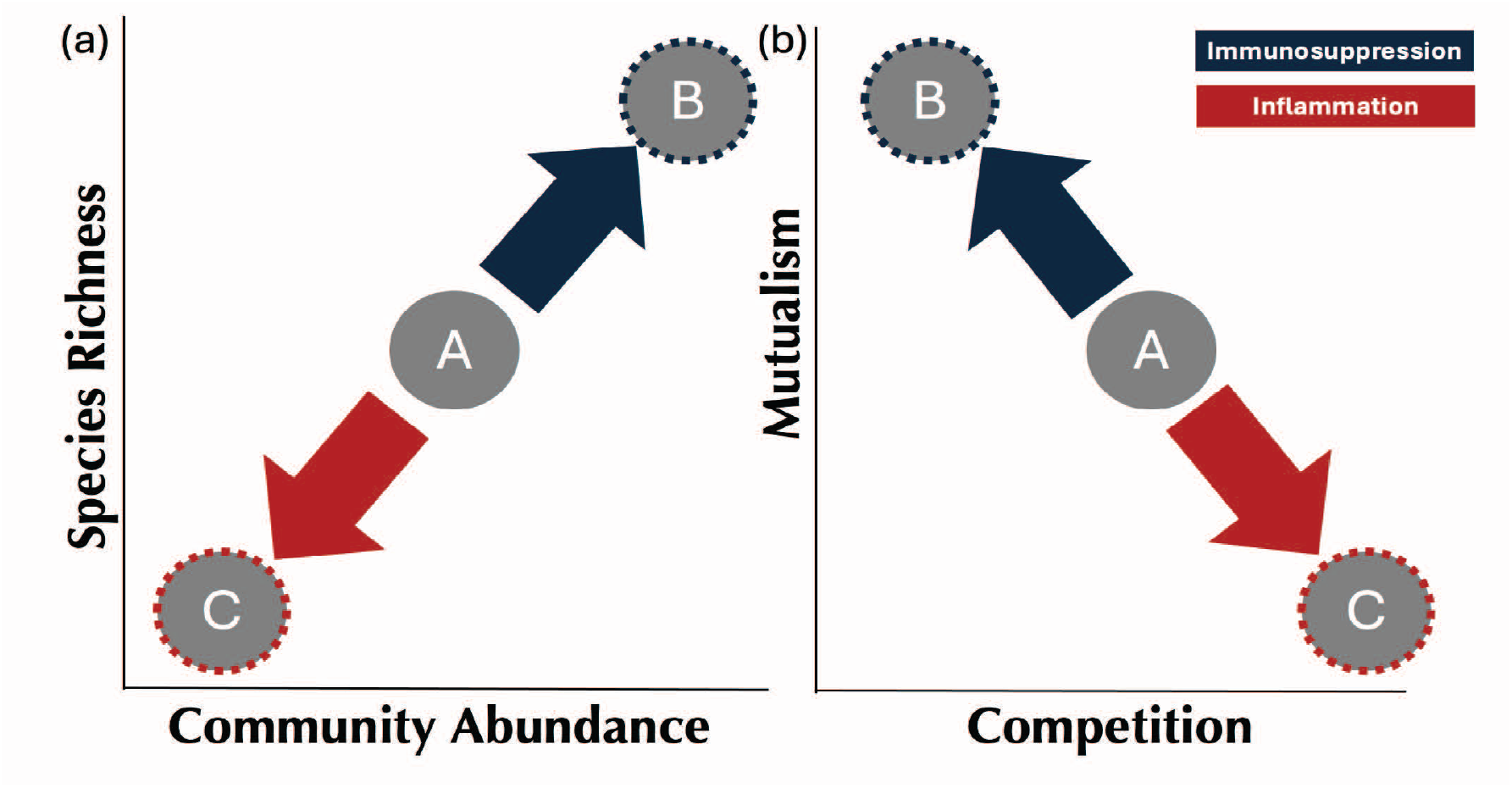
Graphical illustration of our hypothesis for the influence of host immune states on microbial communities. Each circle (A, B, C) represents a microbial community. Panel (a) depicts species richness and total microbial abundance of hypothetical microbiomes, while panel (b) depicts the balance of interactions for the same microbiomes. At host homeostasis, the microbiome is in state A. If host homeostasis is disrupted by inflammation, the microbiome exhibits reduced richness and abundance (shifting toward C), which favors competition. Conversely, immunosuppression increases richness and abundance (shifting toward B), promoting mutualism.

In this paper our goal is to test these theoretical insights using real-world microbiome data. This approach aims to solidify our understanding of the impact of host immune control on the microbial community dynamics. In our paper, we provide a case study analysis as an illustrative example. We demonstrate how both metabolic modeling and genomic analyses can be utilized to elucidate the intricate interplay among microbiome richness, abundance, competition-mutualism interactions, and the host’s immune state.

To quantify the ecological interactions among microbes we use a systems biology approach considering the metabolic “networks of networks” (Muller et al., 2018). Each microbe within a community harbors its own molecular network, made up of genes encoding enzymes that drive different metabolic processes. These metabolic networks interact when two species either can use each others metabolites, or require the same metabolites to grow (Muller et al., 2018). Decoding these interactions requires genomic sequences of the microbes, which are reconstructed into metabolic models. Recent years have seen the development of methods such as ModelSEED (Faria et al., 2023), CarveMe (Machado et al., 2018), and Pathway Tools (Karp et al., 2011) that can generate these networks directly from genome data. This genome-based approach assumes that microbes can execute all metabolic functions for which they possess the corresponding genetic coding (Muller et al., 2018). Once metabolic models are built for all species in a community, their metabolism can be quantitatively analyzed in silico using techniques such as flux balance analysis (FBA) (Orth et al., 2010). Tools such as SMETANA (Species METabolic interaction ANAlysis) (Zelezniak et al., 2015) that are built on the foundation of FBA take this further by calculating metrics to assess competition among microbes by quantifying the overlapping resource use, and cooperation, by assessing the reliance of a microbe on another microbe’s metabolic byproducts (Zelezniak et al., 2015). Unlike methods limited to pairwise interactions, SMETANA evaluates entire communities, predicting how the group functions collectively. This is crucial, as microbial interactions often involve higher-order dynamics beyond simple one-to-one exchanges (Bairey et al., 2016).

We present analyses using The Cancer Genome Atlas (TCGA) dataset (Weinstein et al., 2013), which offers comprehensive information on the genomic, transcriptomic, microbiome, clinical and immune profiles across 33 different cancer types. Investigating microbial ecological patterns in the context of cancer can be insightful as cancer harbors a unique tumor microenvironment characterized with a specific immunophenotype and a distinct microbial community. We examine how the immune response and microbial community characteristics vary across different hosts, we divide cancer microbiome samples into two groups: “high” richness and abundance samples and “low” richness and abundance samples. With these two groups, we then identify the compositional and interaction differences within microbial communities belonging to the two groups as well as differences in host immune features.

## 2 Methods

### 2.1 Dataset

We used the publicly available TCGA pan cancer microbiome dataset from (Narunsky-Haziza et al., 2022; Poore et al., 2020). This dataset comprises bacteria and fungi at both genus and species level. We are using the version of this dataset that has been revised to consider the updated host depletion step to discard any read pairs that mapped to the human genome as described by Narunsky-Haziza et al. (2022). Contaminants has been removed with the help of in-silico methods and literature review ((Narunsky-Haziza et al., 2022). The dataset was particularly useful for our case study as it includes gene expression, clinical and microbiome data across 33 cancer types with more than 17,000 patient samples. In the main text we present our analyses on Stomach cancer (STAD). However; we repeat the analysis across other cancer types which we elaborate more in the supplementary information.

There were 625 RNA seq and 161 WGS STAD samples present in the dataset, all sequenced using Illumina HiSeq. We removed any species across samples whose total count sum was less than 500 reads, and any samples with no species present. We next normalized the OTU table by dividing each species read in a sample by the total reads for that sample. The total reads consisted of reads mapped to human sequences and unmapped reads indicative of microbial sequences. We performed PCA analysis on these tumor samples to determine the presence of batch effects linked to the data submitting center and the experimental strategy (RNA seq or WGS).

### 2.2 Preliminary analyses

#### 2.2.1 Analyzing high and low richness samples

We define the number of OTUs in a given sample as the richness of species observed in that sample. We define the abundance as the sum of the normalized counts of each of the species in a given sample, we take log of that sum to be able to graphically represent the relationship between richness and abundance. To extract samples that show the stark contrast between richness and abundance, we took the first and last quantile samples in each measure. This allows us to capture key ecological differences between high and low richness and/or abundance microbiomes. As a control, we also analyzed samples arbitrarily and randomly assigned to be “high” or “low.”

#### 2.2.2 Community composition analysis among high and low groups

We first plotted the rank abundance curves for each sample. A rank abundance curve is the normalized abundances of each OTU in a sample, ordered by their abundance, going from high abundance to low. We next determined the average abundance at each rank for samples in both the low and high groups. To determine differentially abundant species across high and low groups, we used the Analysis of Composition of Microbiomes (ANCOM) statistical (Mandal et al., 2015) method. This methodology can effectively deal with the compositional nature of the microbiome data. We considered additional covariates such as age of diagnosis, gender and the data submitting center for the ANCOM model.

### 2.3 Mutualism and Competition Interaction Landscape between Groups

#### 2.3.1 Determining co-occurring communities

We first filtered the OTU table to only consider species that are present in more than 50% of the samples. This allows us to only retain species that are more likely to play a biologically relevant role in the community. We then used the filtered OTU table to construct a correlation matrix to determine which species tend to covary together across samples. To do that, we conducted Spearman correlation analysis to determine the pairwise correlation among species present in the samples. To construct the co-occurance network, we considered species whose pairwise correlation coefficients are greater than 0.6 in absolute value and Benjamini-Hochberg adjusted p value is less than 0.01. We then used this correlation matrix as an input to the fast greedy community detection algorithm (Clauset et al., 2004) to determine co-occurring communities within the samples. The algorithm treats each species as a separate node and then iteratively joins them to maximize the modularity of the network. This algorithm identified co-occurring sub-communities within both the low and high groups. We only consider a sub community that consists of more than 10 species. Subsequently, we evaluate the collective metabolic potential of each co-occurring sub-community (see section 2.3.2). We next compute the average metabolic potential across all sub-co-occurring communities identified within each group.

#### 2.3.2 Determining metabolic interactions

For each of the species present in the subcommunity we download a reference genome from the NCBI database. We used the species genome files to reconstruct a genome-scale metabolic model using a python package CarvMe (Machado et al., 2018). The CarvMe package allows fast automated reconstruction of genome-scale metabolic models for microbial species. For our main analyses, we used the default option to predict the uptake and secretion capabilities of an organism from the genetic evidence only, it yields a simulation ready model without gap filling for any particular media. However, media can be provided as an additional input for reconstruction. To check the robustness of our results, we also created metabolic models with gap filling for M9 and LB experimentally verified media. We considered gap filling the model by media to assess if the media itself can greatly influence the overall metabolic potential of the subcommunity. Our results remained qualitatively unchanged. Once we had the metabolic files ready for each species in the subcommunity, we then used them as an input to the SMETANA algorithm developed by (Zelezniak et al., 2015) to determine the metabolic resource overlap (competition) and mutualistic interaction potential (mutualism) for that subcommunity.

We next assessed the metabolic potential of a single species in relation with the other members in the community. We specifically focused on the metabolic interaction potential of differentially enriched species that we identified using ANCOM observed in the low and high richness and/or abundance samples. This can reveal intricate metabolic dependencies and synergistic relationships, shedding light on how specific species contribute towards community characteristics such as abundance. Understanding these interactions can help us pinpoint keystone species that play a disproportionate role in governing community dynamics.

To determine the metabolic interaction potential of the species that were differentially abundant in the high and low richness and/or abundance samples, we conducted a series of in-silico simulations with a varying number of other species. Specifically we examined the competition and mutualism score from the SMETANA tool when considering the interaction between the differentially enriched chosen species with a number of randomly selected species present within the same community. The interaction pairings are based on the available number of species present in the community, where pairings with the chosen species can consist of 4, 6, or 8 species. Each interaction scenario was repeated at least 10 times to robustly determine the interaction. We also conducted the same in-silico analysis using randomly selected species, excluding the chosen species to be able to establish a baseline comparison for competition and mutualism scores within that sub community.

### 2.4 Determining differentially enriched immune genes

We downloaded the gene expression data available for the STAD samples. We conducted DEseq analysis on the expression counts of immune genes to ascertain significant differences between the high and low groups.

We obtained raw gene expression counts for the STAD samples. We narrowed down our genes list to only immune genes with the help of the nCounter PanCancer Immune Profiling Panel (Cesano, 2015). It comprises 770 immune markers that cover both the adaptive and the innate immune response (B cells, T cells, TH1 cells, Treg, CD45, CD8, Cytotoxic cells, Dendritic cells, Macrophages, Mast cells, Neutrophils, and Natural Killer cells). We filtered out low read counts, retaining genes with counts per Million (CPM) > 1 in at least 50% of samples, and normalized gene expression counts across samples.

We then used the normalized gene counts table to perform differential gene expression analysis across both low and high groups using the DEseq package in R. We selected for immune genes enriched in both high and low groups that had a fdr corrected p adjusted value lower than 0.05, and the absolute log2 fold change to be greater than 0.5.

### 2.5 Survival analysis between groups

We performed survival analysis to determine if our categorization based on either species richness, abundance or both had a prognostic difference in the survival status of the patients between the low and the high groups. We generated a Kaplan-Meier survival curve to visualize the survival probabilities for low and high group. Additionally, a log-rank test was employed to compare survival curves between groups.

## 3 Results

Using the microbiome data obtained from the TCGA STAD tumor tissues, we aimed to investigate the impact of variations in microbial community characteristics, including community abundance and richness, on the ecological interactions observed within the community and their association with the immune response. When plotting community richness and abundance observed in each of the samples (Figure 2a) we saw an overall increase in abundance with an increase in richness. We next clustered the samples into low and high groups using quantile based clustering along both the richness and the abundance axis. In the subsequent text we use term the low and high groups, as observed in (Figure 2a), comprising of samples with low richness and abundance and high richness and abundance respectively.

**Figure 2:**
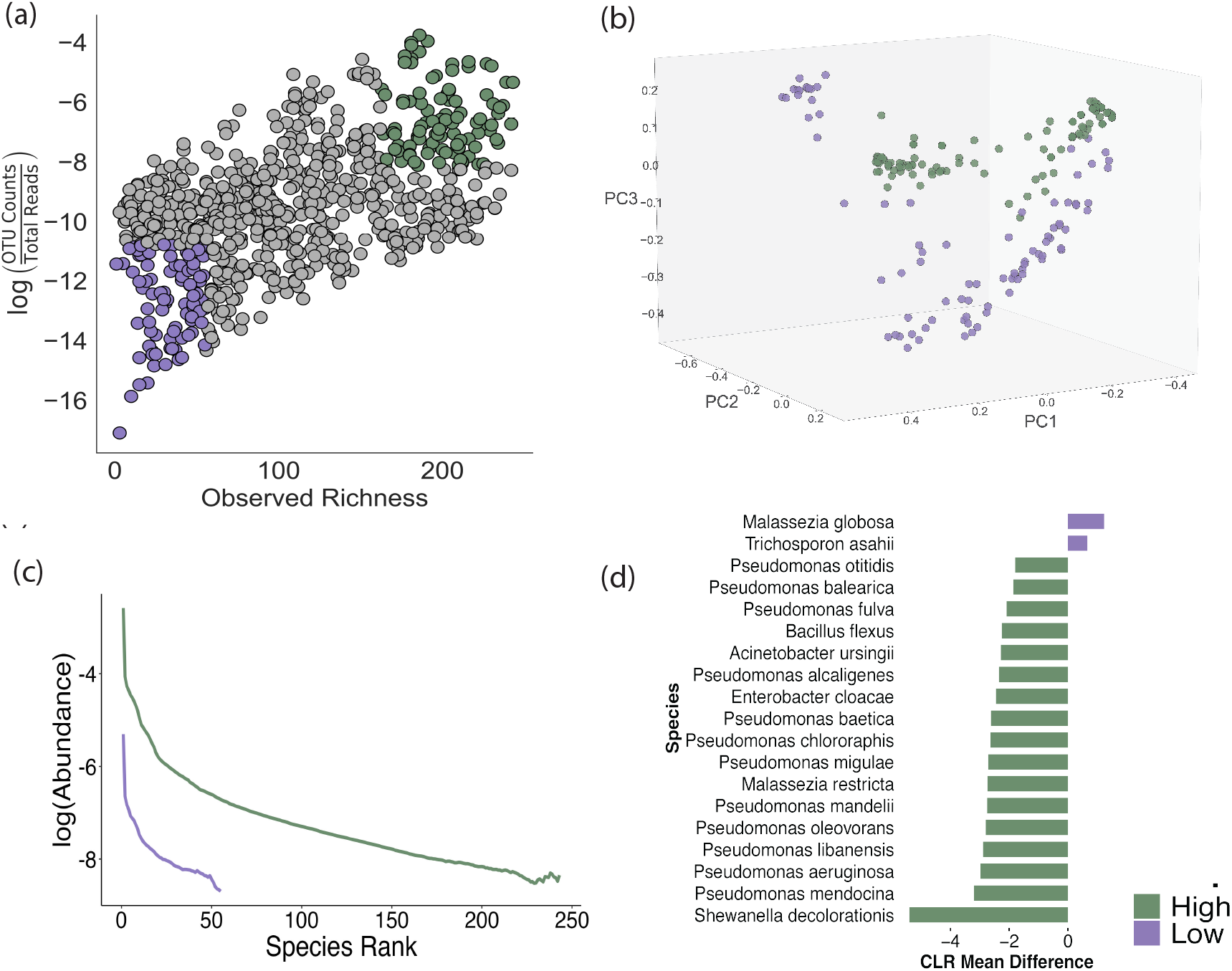
Analyzing differences in composition between cancer microbiomes with high and low richness. **a)** Samples were categorized into high and low groups using quantile-based clustering. Samples from the lower and upper quantiles were assigned to the low and high groups, respectively, across both the x-axis. **b)** PCA analysis on microbiome samples across low and high group. **c)** Each sample’s rank abundance curve is depicted, with bold lines indicating group averages. **d)** Significantly differentially abundant species between low and high groups, as determined by the ANCOM model.

The two groups exhibited distinct rank abundance curves, as these curves provide insights into the evenness of the microbial community. Evenness describes the distribution of community members in terms of their abundance, indicating whether certain species are dominant while others are rare, or if species abundances are more evenly distributed. In both the groups, species abundances were evenly distributed with no stark contrast in the slope of the curve among the two groups (Figure 2c).

We next identified differences in the composition of the microbiome among the two groups. We observed a notable increase in the abundance of the phylum Proteobacteria in the high group compared to the samples categorized under the low group (SI Figure 2a). Upon examination at the genus resolution (SI Figure 2b), we noted a general rise in Shewanella in the high group (SI Figure 2.7b). Upon performing PCA, we observed that samples cluster based on group membership (Figure 2b). We also observed species from genus Shewanella being differentially enriched among the low and the high group using ANCOM which is a more robust statistical assessment of quantitative differences in composition among groups (Figure 2d). Binning data for analysis can introduce bias, to address this limitation, we randomly selected 500 sets of high and low groups from the data. This approach allowed us to assess the robustness of our findings across different group assignments and mitigate potential biases associated with binning. We found that for 73% of cases, we observed no significant enrichment of species in either the low or high group based on ANCOM analysis (SI Figure 5b). Among the remaining 27% of cases where species were significant, the genera Malassezia, Shewanella, and Pseudomonas were most frequently identified as significant (SI Figure 5c).

Next, we evaluated the differences in metabolically mediated interactions between the two groups. We initially identified co-occurring sub-communities within both the low and high groups. In the low group, there were two cooccurring sub communities (SI Figure 3a,c). In the high group, there were fifteen co-occurring sub communities detected (SI Figure 3b,d). When comparing metabolically mediated interactions among the low and high groups, the high group exhibited on average higher level of mutualism compared to the low group. With mutualism showing a more pronounced average change compared to competition (Figure 3a). These insights remained robust when determining metabolically mediated interactions across M9 and LB media (SI Figure 4). Overall across all co-occurring sub communities for both low and high groups, we observed both species richness and abundance to increase with mutualism and decrease with competition. Where the linear regression model between species richness and mutualism exhibited a significant p value and the highest *R*^2^ value among all the four models that we tested (Figure 3b). One interesting exception to this pattern was sub-community #8 observed in the high group (SI Figure 3d). Despite having comparable levels of competition and mutualism to the communities in the low group, this sub-community displayed a considerably higher level of abundance compared to the low group sub-communities (SI Figure 3c). Sub community #8’s abundance is reflective of the presence of *Shewanella decolorationis*. As previously noted, the high group exhibits a higher relative abundance of *S. decolorationis*, with this species exclusively present in sub-community #8.

**Figure 3:**
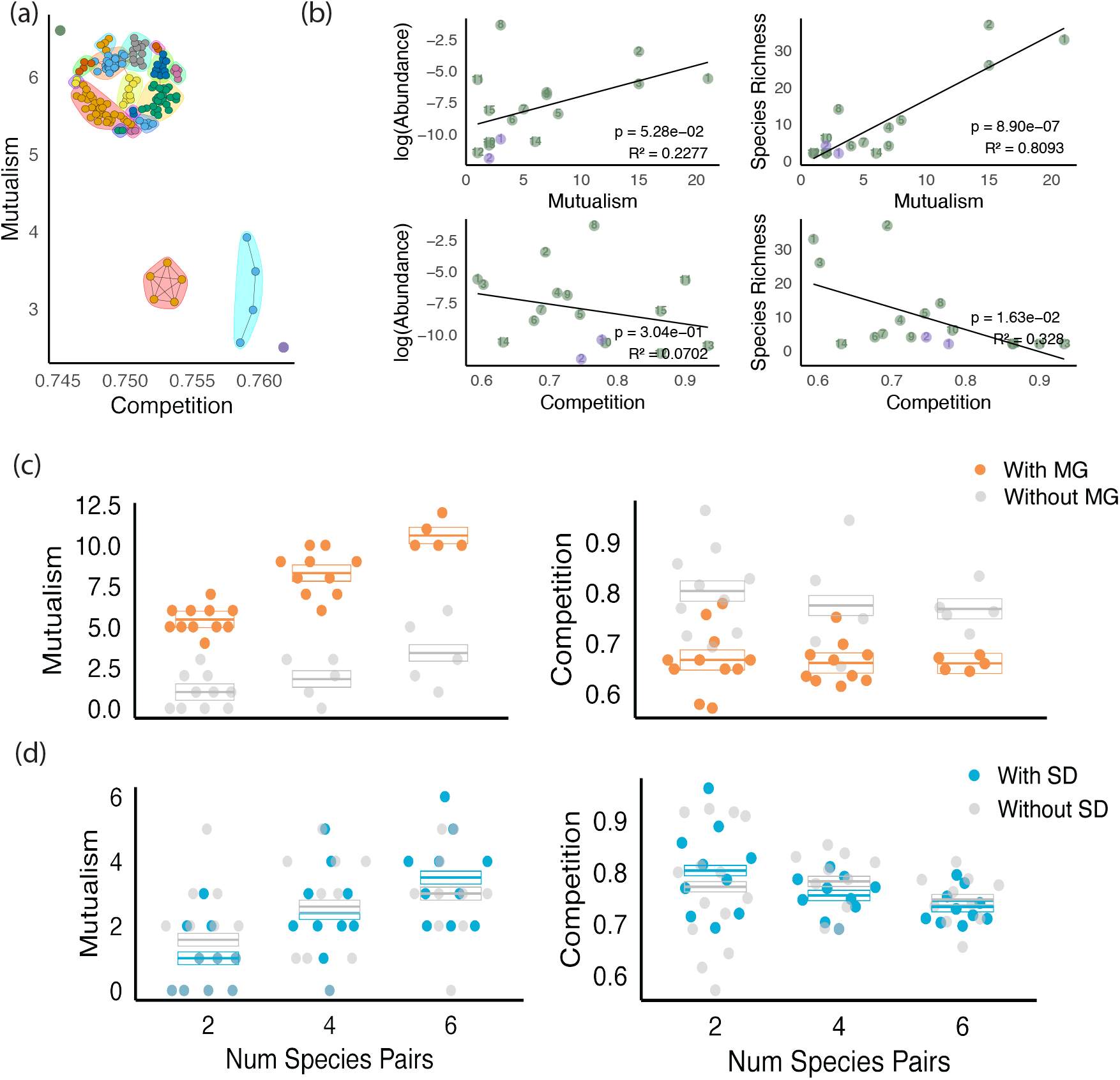
**(a)** Represents the average competition and mutualism observed across all the co-occurring sub communities identified within both high and low samples as indicated by the networks. Each node in the network represents a species, and the lines connecting nodes indicate either positive (black lines) or negative correlation (red lines). Sub-communities are identified by different colored shading. **(b)** Represents the linear regression analysis between mutualism and competition scores and species richness and abundance all for subcommunities observed within both low and high groups. Each dot is labeled by the sub community id. **(c-d)** Portrays the interaction potential of differentially enriched species Shewanella decolorationis, and Malassezia globosa

To determine the individual metabolic interaction potential of the most differentially enriched species across the two groups. Using members of subcommunity #8, we simulated metabolic models of microbial community both with and without S. decolorationis. We found that *S. decolorationis* does not significantly alter the level of mutualistic or competitive interactions among community members compared to interactions observed in within the same sub community lacking *S. decolorationis* (Figure 3d). This shows that despite the elevated abundance of sub-community (#8) relative to those in the low group, *S. decolorationis* appears to have minimal impact on the competitive or mutualistic dynamics within the community.

We similarly assessed the metabolic interaction potential of *Malassezia globosa*, the species most differentially enriched in the low group. Here, we did observe a notable effect (Figure 3c): *M. globosa* appears to enhance mutualism within the community while reducing competition among species within the same sub-community, compared to interactions observed in within the same sub-community in absence of *M. globosa* (Figure 3c).

We then evaluated whether there were any notable differences in immune response observed between the low and high groups. We observed significant immune markers differentially enriched across groups (Figure 4). We find *HLA-DQA1* to be significantly enriched in low group whereas *C8G, MASP1, PLA2G1B*, and *TNFRSF17* enriched in the high group (Figure 4).

**Figure 4:**
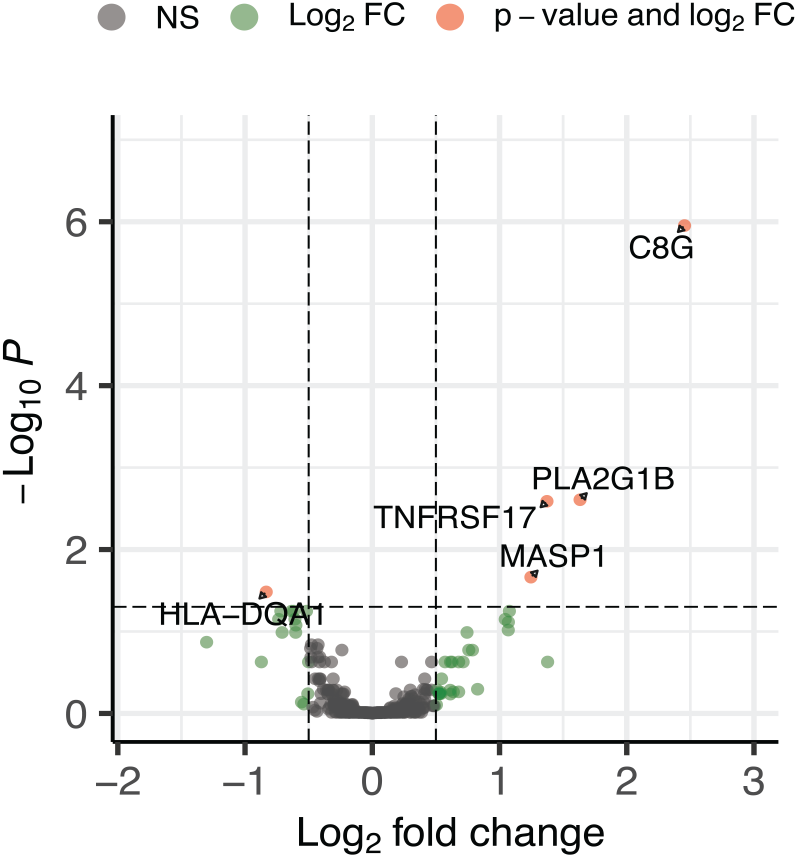
Depicts the volcano plot showcasing differentially enriched immune markers between the high and low groups. A positive fold change signifies immune markers enriched in the high group, while a negative log fold change denotes immune markers enriched in the low group.

Finally, we asked if the ecological properties of the microbiome is correlated with the outcome of the cancer. We detected a significant difference in survival rates between the low and high groups, with patients in the high group exhibiting a poorer prognosis (Figure 5a, see SI for more information on survival analysis). To address potential biases in group assignments affecting the observed survival differences, we randomly assigned samples to either the low or high group 500 times and repeated the survival analyses. We found that only about 5% of these random groupings produced a statistically significant survival difference between the two groups, demonstrating that our results are statistically robust across different group assignments (SI Figure 5a). Furthermore, we found that patients who retained their tumor status had a negative odds ratio correlated with the abundance of the microbial community, compared to those who transitioned to a tumor-free state. Interestingly, species richness did not show a significant impact on the tumor status of the patients (Figure 5b).

**Figure 5:**
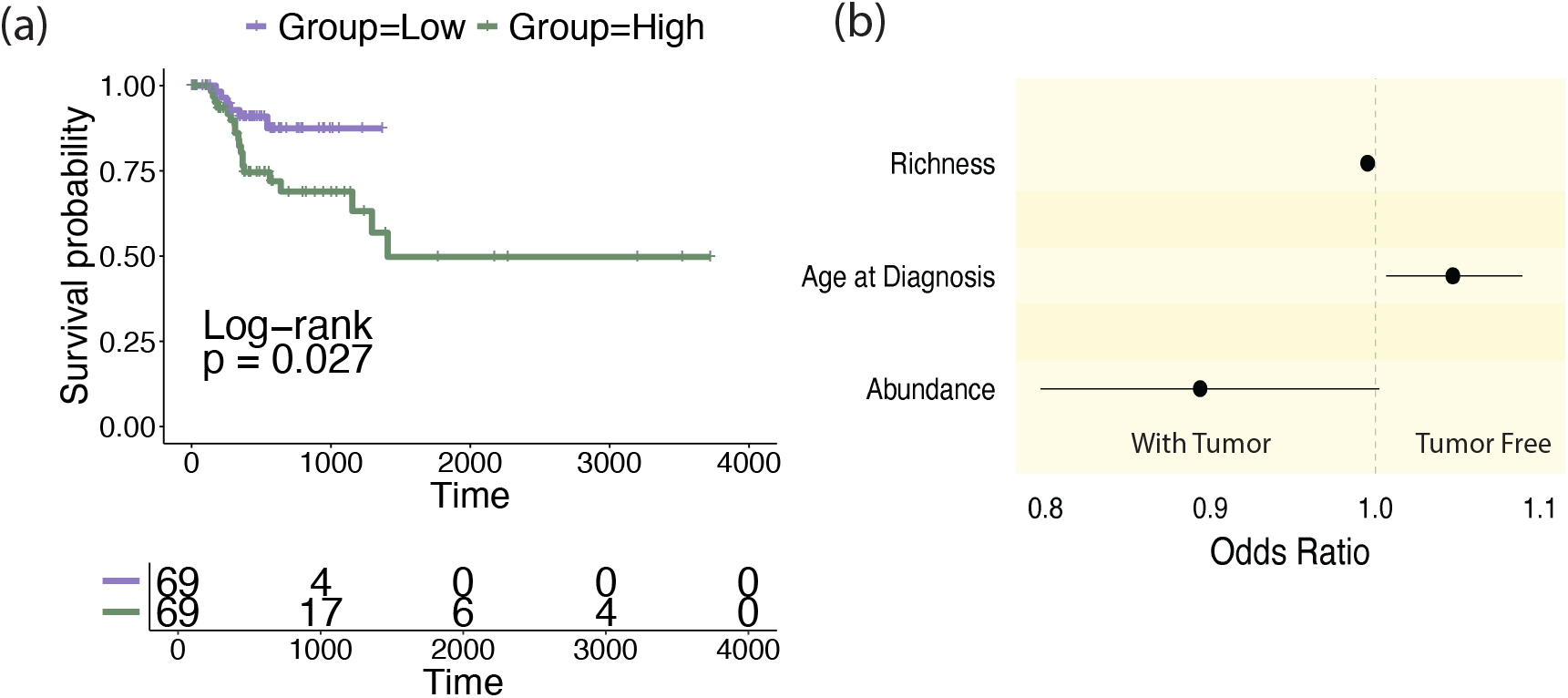
**(a)** Kaplan-Meier survival curves are presented for the low and high groups, with p-values from the log-rank test indicating statistical significance. **(b)** Represents an odds ratio plot across species richness, age of diagnosis, and community abundance for patients who had tumors or those who became tumor-free.

## 4 Discussion

In this study, we used the TCGA stomach cancer dataset as a case study to illustrate how community ecology principles can inform microbiome studies in health context. Our results illustrate the type of analyses that can be conducted with additional datasets to connect microbiome composition, its ecology in terms of metabolically mediated interactions among community members and the microbiome’s connection to the immune response.

We first establish the relationship between microbiome community richness and abundance. Both of these metrics are important measures of community fitness. While microbiome species richness has historically received significant attention (Bäckhed et al., 2012; Le Chatelier et al., 2013), the importance of abundance as a biomarker for health status of the microbiome is often overlooked. These metrics can be correlated, jointly informing community fitness, or they may act independently, driving distinct community dynamics. Within stomach cancer samples, we observe a positive linear correlation between microbiome richness and abundance. We show that just by simply clustering samples into low and high groups based on richness and abundance we are able to highlight the combined prognostic role of these metrics for patient survival. Moreover, our findings indicate that abundance, rather than richness, is linked with a increased likelihood of tumor presence in patients, underscoring the significance of abundance as a prognostic indicator. When we plot additional cancer types, such as Ovarian and Heck and Neck Sarcomas, across richness and abundance we find additional instances where richness and abundance are correlated (SI Figure 6). This suggests that the microbial ecologies of tumor samples from these cancer types may be shared from that of STAD, and the prognostic role of the two fitness metrics may align similarly.

We further demonstrate that differences in community characteristics correspond to a distinct ecological interaction landscapes. We specifically quantified ecological interactions as metabolically mediated competition and cooperation, inferred by geneme-based metabolic modeling. Our findings reveal that samples characterized by high levels of abundance and richness exhibit significantly higher levels of mutualism, indicated by the greater sharing of metabolites among members. Interestingly we observe that an increase in diversity within the microbiota does not correspond to an increase in competition among species but in turn allows for an increase in cross-feeding interactions. On average, we observe greater community abundance for highly diverse co-occurring communities. These insights are consistent with our theoretical findings that conclude a more diverse community with high community abundance observed for microbial communities that interact highly mutualistically (Abbasi & Akçay, 2022). Several studies have revealed a positive stabilizing effect of mutualisms in promoting species coexistence. A high level of mutualism among community members renders the community to be self sufficient to perturbations in the dynamic environment (Culp & Goodman, 2023). However our analysis did reveal an exception to the above mentioned trends, where we observed that co-occurring communities despite harboring a higher community abundance were species poor and observed an elevated competition among the community members. It was driven by the taxa (*S. decolorationis*) whose role in the cancer microbiome remain unresolved.

In a healthy host, the stomach is known to support a low microbial community abundance in comparison to microbiome abundance observed in the small and large intestine (Sender et al., 2016). The cap on the microbial community abundance is sustained as a result of interplay of several host filters including the low pH environment (Williams, 2001), and the immune cell population.

However in a perturbed state such as cancer there may be an increase in microbial community abundance that can signify dysbiosis and an altered immune control on the microbiome (Zheng et al., 2020). Our results support these insights as we observe poor prognosis for patients that have high abundance and richness and are less likely to regress into a tumor free state. We hypothesize that the samples in the low group may have an intact immune response in comparison to the high group samples. A hallmark of a functioning immune response is ability for the host to undergo immune surveillance that entails the recognition of antigens. Consistent with this, we find that *HLA-DQA1*, whose primary function is in antigen processing and presentation, exhibited the greatest positive fold change in expression in the low group. Previously, *HLA-DQA1* was shown to be significantly associated with long term survival for patients with soft tissue sarcomas (Bae et al., 2020), and its absence led to an increased incidence of gastritis (Azuma et al., 1998).

Conversely, in the high group we find the immune-associated genes *C8G, MASP1, PLA2G1B*, and *TNFRSF17* to be significantly enriched. *C8G* is a component of the complement system which plays a role in defense against pathogens (Schreck et al., 2000). There is limited insight into the role of *C8G* in STAD, however, its prognostic significance has been underscored in the context of colorectal cancer (CRC). In a study by Zhao et al. (2022), *C8G* was identified as one of the eight immune genes contributing to the overall survival of CRC patients. Elevated expression of *C8G* was observed in the high-risk group, which exhibited poorer patient survival outcomes compared to the low-risk group. *MASP1* is known to play an active immunological role in STAD, although the expression of *MASP1* is known to be reduced in tumor state in comparison to normal stomach tissues, gene ontology analysis revealed that *MASP1* is linked to pathways that play a role in mismatch repair and gastric cancer (Zhang et al., 2022). *PLA2G1B*, or pancreatic group IB secreted phospholipase A2, plays a crucial role in the digestion of dietary and biliary phospholipids within the digestive tract (Hui, 2019). Where inhibiting *PLA2G1B* improves overall metabolic health in the digestive track, and lowers the risk of developing obesity, diabetes and liver diseases (Hui, 2019). Moreover reduced activity of *PLA2G1B* is associated with a reduced risk of colorectal cancer (Hui, 2019) in humans and experimental coloitis in mice (Haller et al., 2023). *TNFRSF17*, Tumor Necrosis Factor Receptor Superfamily Member 17, is known to be involved in B cell homeostasis. Guan et al., 2021 identified *TNFRSF17* as one of the prognosis-associated immunity genes which was observed to be associated with poor prognosis for patients with gastric cancer. Our analysis revealed immune markers, which were highly enriched in the high richness and abundance group, to be associated with poor cancer prognosis. Hence indicating a perturbed immune state in the high richness and abundance group. This underscores the importance of integrating community ecology principles into biomarker discovery.

Our results do not capture the comprehensive landscape of ecological interactions as observed within the microbiome. More specifically we only examine passive competition where species compete with each other via resource consumption, we acknowledge that microbes can also actively compete with each other, engaging in chemical warfare which can also affect community dynamics (Ghoul & Mitri, 2016). Furthermore, we only consider the changes in immune gene expression among the low and the high groups. Further analysis can focus on exploring the variation in specific immune gene expression in relation to microbiome communities or individual microbial species. Such investigations, although not within the scope of this study, have already been undertaken in studies such as (Huang et al., 2023).

Our results underscore the utility of community ecology principles in understanding microbiome characteristics and their impact on human health. Our analyses demonstrated that microbiome richness and abundance can serve as important biomarkers influencing patient prognosis and disease progression. We showed that metabolically mediated interactions, particularly competition and mutualism, can be informative about the microbial richness and abundance community characteristics. Additionally, by incorporating immune context when analyzing the microbiome, we gained insight into the strength of the host immune leash on the microbiome. These results confirm previous theoretical predictions and can help further our understanding of the collective role of microbial interactions and the immune response in informing microbiome richness and abundance. We hope our study inspires similar analyses on additional datasets to further explore the role of community ecology and immune response in shaping the microbiome.

## 5 Code Availability

The analysis was implemented in Python. The code is available at GitHub: https://github.com/EemanAbbasi/TCGA_Cancer_Microbiome_Analysis_Pipeline

## 7 Author Contributions

All authors designed the study, Eeman Abbasi implemented the analysis. All authors contributed to writing the manuscript. All authors gave the final approval for publication.

## 9 Supplementary Information

### 9.0.1 Survival Analysis

In our TCGA survival analysis, we’re comparing disease-specific survival (DSS) between “Low” and “High” microbiome groups. We also take into account the status variable in our dataset. The variable indicates whether a patient experienced the event (death): status = TRUE: The patient died within the observed period. status = FALSE: The patient was censored (alive, lost to follow-up, or study ended without death). Censored patients contribute to the survival curve and number at risk but don’t add to the event count for statistical comparison. For our data below is the distribution of the status variable among the two groups.

Where:

| Group | False | True |
| --- | --- | --- |
| High | 50 | 19 |
| Low | 63 | 6 |

#### High Group

19 deaths (TRUE), 50 censored (FALSE), total 69 patients (27.5% event rate).

#### Low Group

Only 6 deaths (TRUE), 63 censored (FALSE), total 69 patients (8.7% event rate).

The zeros in the risk table for the “Low” group occur because, by this time all 69 “Low” group patients are either:

#### Dead

(6 patients, status = TRUE, events occurred during the timeframe).

#### Censored

(63 patients, status = FALSE, likely censored due to being alive or lost to follow-up).

The risk table shows the number of patients at risk (those not yet dead or censored) at each time point. For the “Low” group, the low event rate (6 deaths) and high censoring (63 patients) mean that the number at risk drops rapidly. If most of the 63 censored patients are censored before 1600 days (e.g., median censoring time 800–1000 days), the “Low” group has no patients left at risk after 1600 days, resulting in zeros in the risk table.

### 9.0.2 SI Figures

**Figure SI1:**
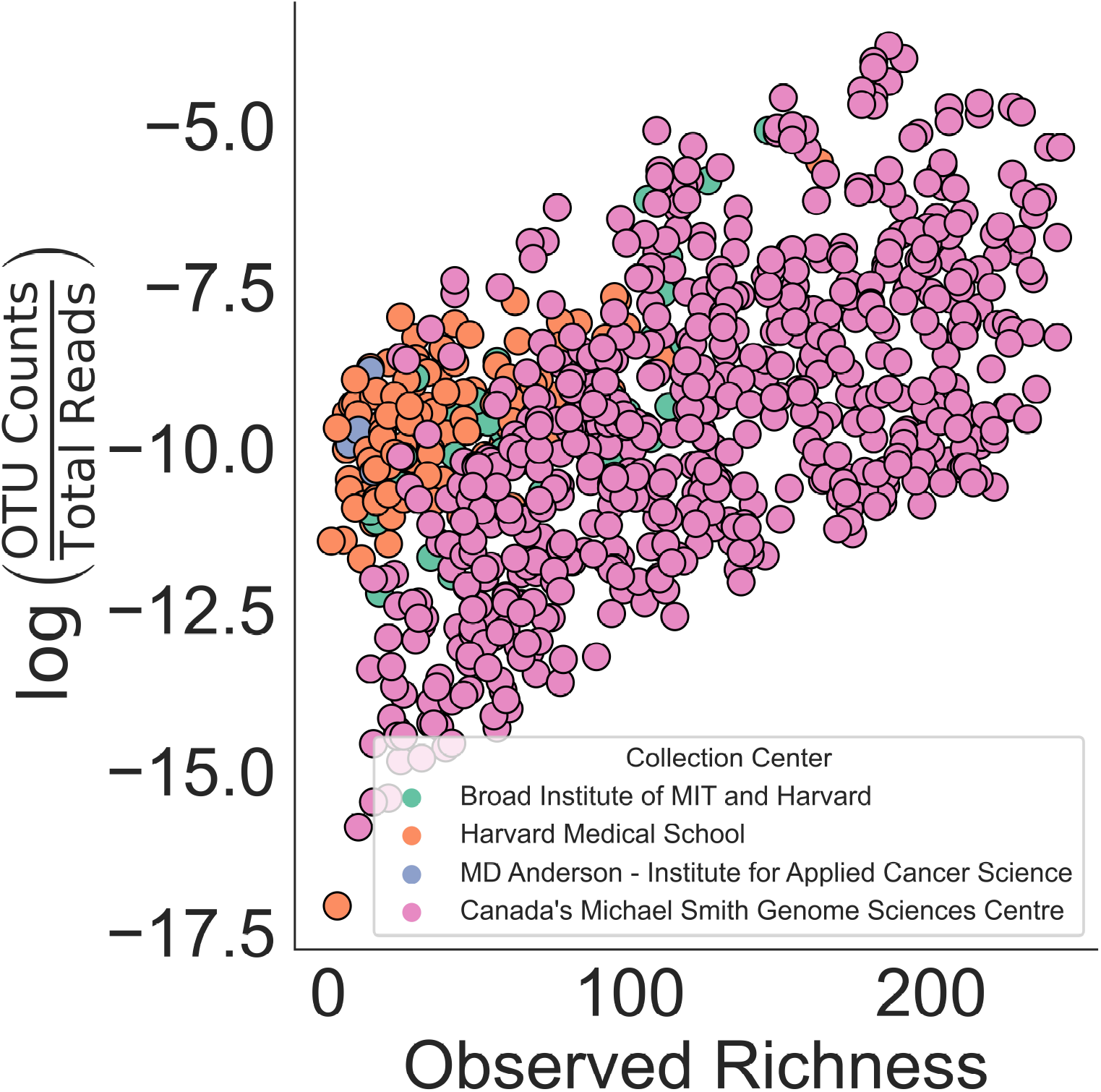
(a) Samples are annotated based on their data submitting center.

**Figure SI2:**
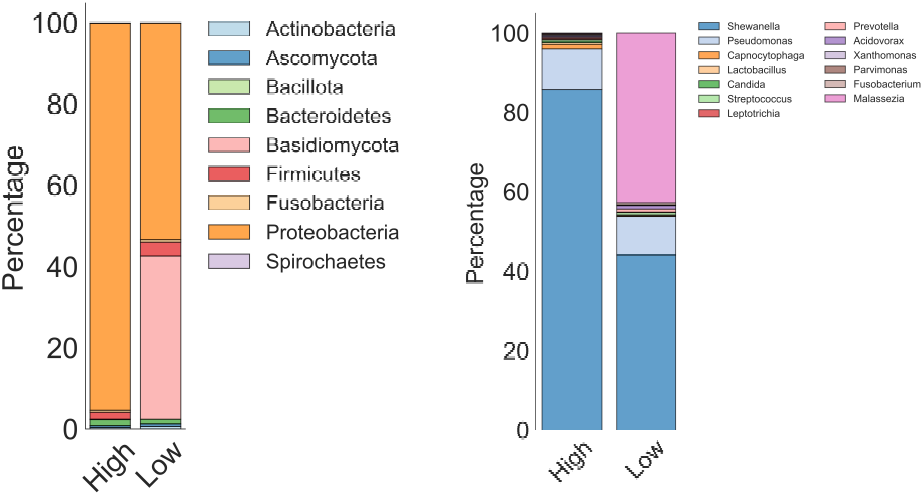
Relative abundance differences observed at the a) Phylum and b) Genus level across groups (top 13 genera shown).

**Figure SI3:**
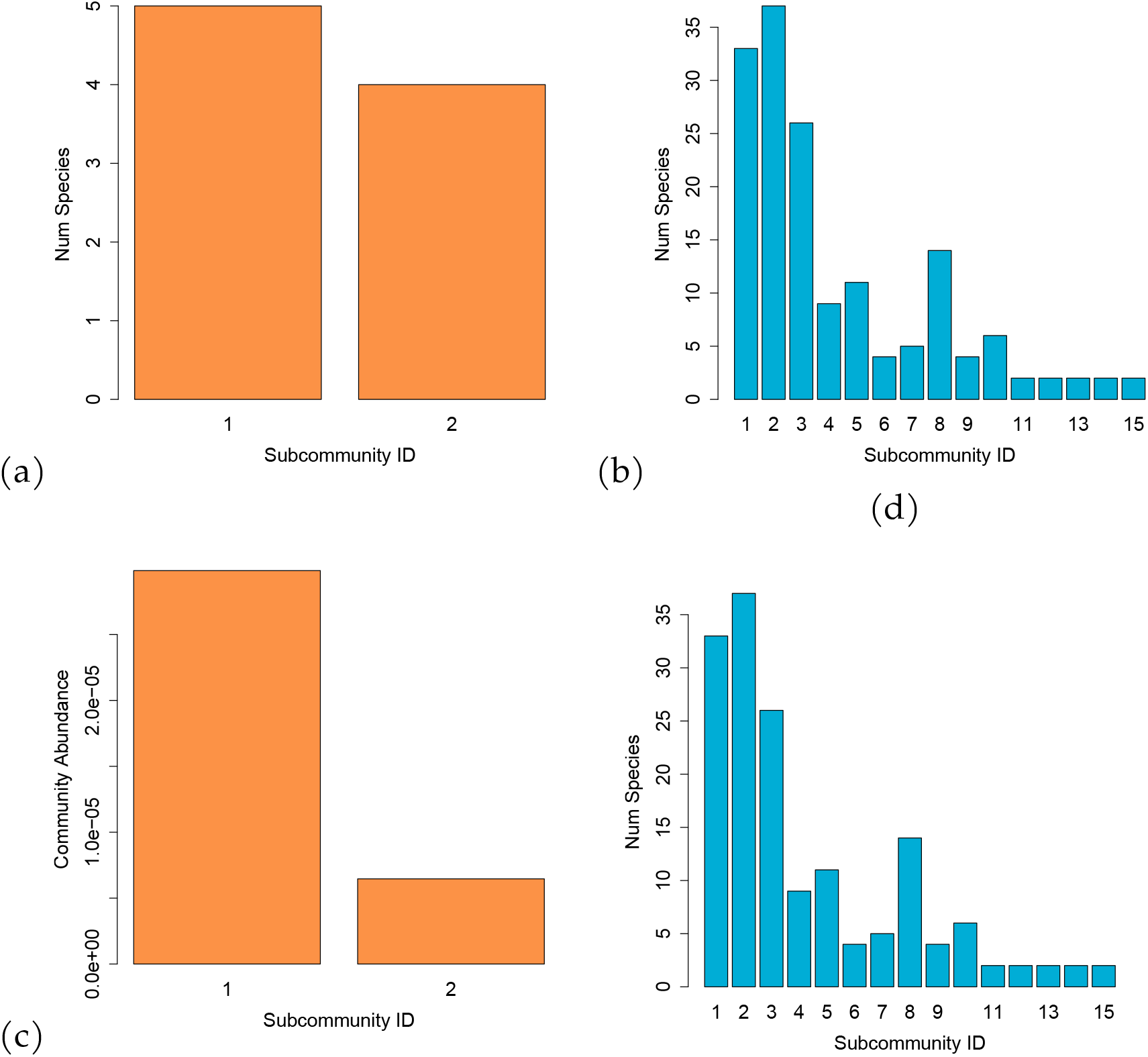
Illustration of co-occurring sub-communities across low and high groups, with orange bars representing sub-communities in the low group and blue bars representing those in the high group.

**Figure SI4:**
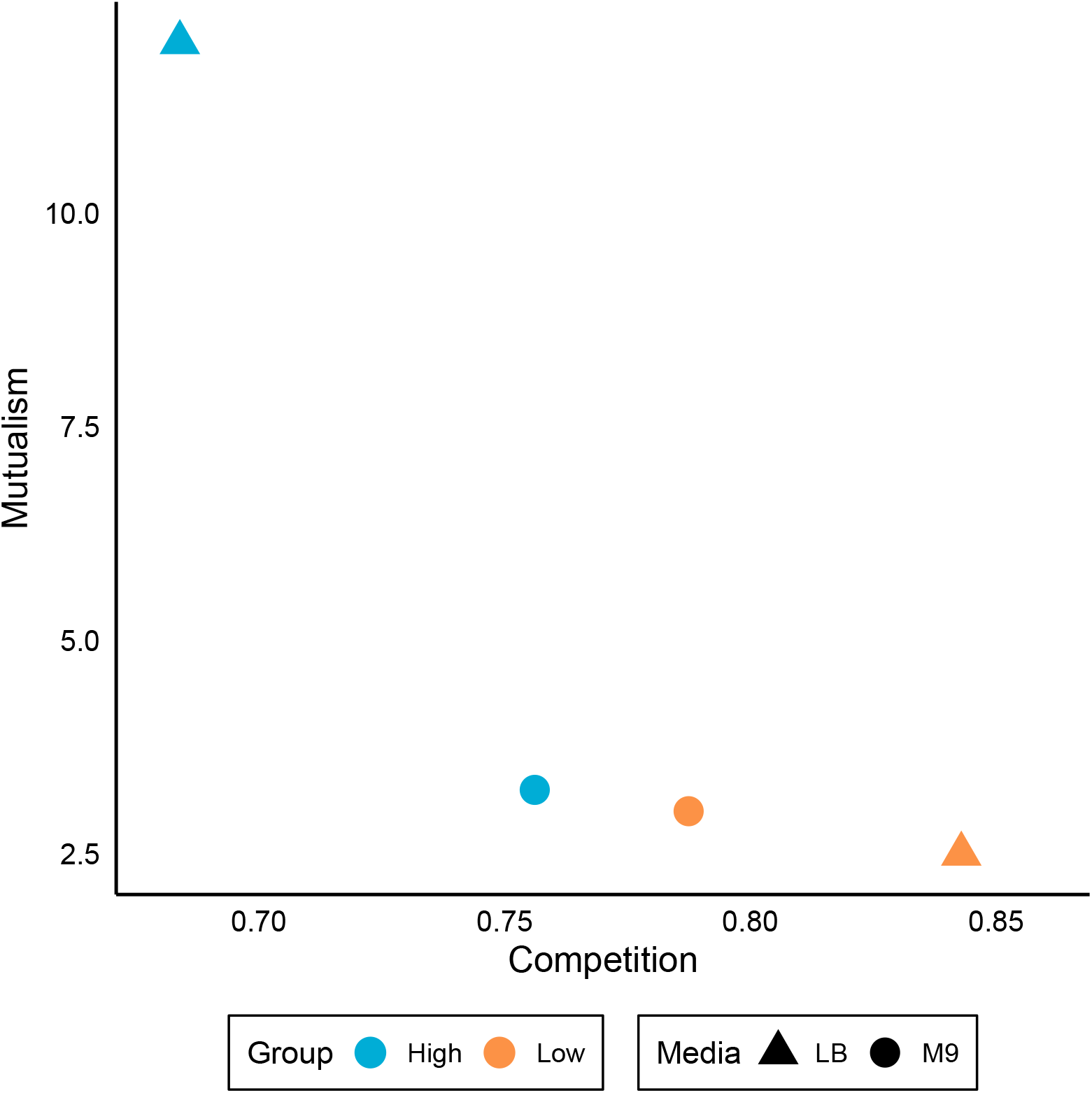
Average mutualism and competition scores for M9 and LB media across both high and low group.

**Figure SI5:**
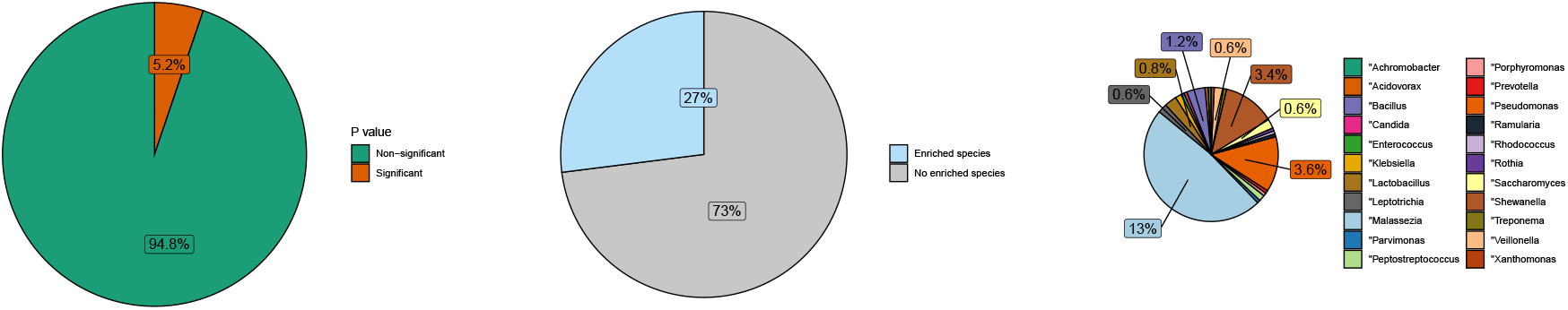
The samples were randomly split into low and high groups, and this partitioning was repeated across 500 simulations. The figure displays the summary outcomes of the analysis. **(a)** Depicts the percentage of statistically significant p-values indicating survival differences between the low and high groups. **(b)** Illustrates the percentage of enriched species and no significant enriched species observed during ANCOM analysis. **(c)** Shows the percentage distribution of enriched genera observed in part b, with all unmarked genera having a percentage of 0.2%. Note, this pie chart only adds up to 27%.

**Figure SI6:**
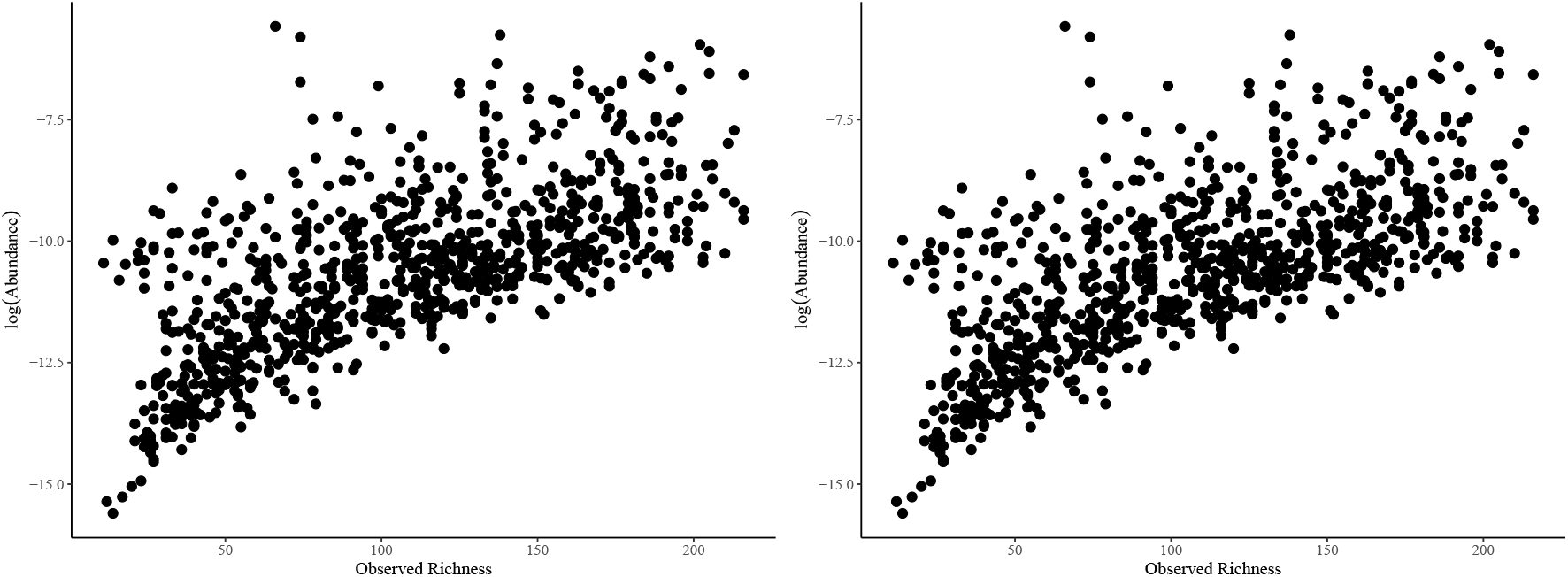
(a) Ovarian, (b) Head and Neck Sarcoma TCGA samples when plotted across both richness and abundance.

